# Playing Musical Chairs in Big Data to Reveal Variables’ Associations

**DOI:** 10.1101/057190

**Authors:** Hugues Aschard, Bjarni Vilhjalmsson, Chirag Patel, David Skurnik, Jimmy Yu, Brian Wolpin, Peter Kraft, Noah Zaitlen

**Author notes:** These authors contributed equally (Peter Kraft, Noah Zaitlen).

## Abstract

Testing for associations in big data faces the problem of multiple comparisons, with true signals buried inside the noise of all associations queried. This is particularly true in genetic association studies where a substantial proportion of the variation of human phenotypes is driven by numerous genetic variants of small effect. The current strategy to improve power to identify these weak associations consists of applying standard marginal statistical approaches and increasing study sample sizes. While successful, this approach does not leverage the environmental and genetic factors shared between the multiple phenotypes collected in contemporary cohorts. Here we develop a method that improves the power of detecting associations when a large number of correlated variables have been measured on the same samples. Our analyses over real and simulated data provide direct support that large sets of correlated variables can be leveraged to achieve dramatic increases in statistical power equivalent to a two or even three folds increase in sample size.

## Background

Performing agnostic searches for association between pairs of variables in large-scale data, using either common statistical techniques or more complex machine learning algorithms, faces the problem of multiple comparisons. This is particularly true for genetic association studies, where contemporary cohorts have access to millions of genetic variants as well as a broad range of clinical factors and biomarkers for each individual. With billions of candidate associations, the identification of a true association of small magnitude is extremely challenging because such signals compete with the large number of correlations that will emerge by chance. The standard analysis approach currently consists of looking at the data in one dimension (i.e. testing a single outcome with each of the millions of candidate genetic predictors) and applying univariate statistical tests-the commonly named GWAS (genome-wide association study) approach^1,2^. To increase power, GWAS rely on increasing sample size in order to reach the very stringent significance level that accounts for the multiple comparisons. The largest studies to date, including hundreds of thousands of individuals across dozens of studies, have been pushing the limit of detectable effect size^3,4^. For example, researchers are now reporting genetic variants explaining less than 0.01% of the total variation of body mass index (BMI)^4,5^.

Apart from the cost of genotyping hundreds of thousands of cases and controls, this brute force approach has practical limits. Sample size cannot be increased indefinitely, especially for rare diseases and diseases for which there is no registry. More importantly, this approach does not leverage the large amount of additional phenotypic and genomic information measured in many studies^6–8^. A commonly discussed alternative to improve statistical power consists of applying multivariate analyses that combine tests of multiple phenotypes with one (or multiple) predictors of interest^9-12^. Standard multivariate analysis, although they offer a gain in power, have two major drawbacks. First, because they are built on a composite null hypothesis, a significant result can only be interpreted as an association with any one of the predictors. While this is useful information for screening purposes, it is insufficient when the ultimate goal is to identify specific genotype-phenotype associations. Moreover, it makes the replication process more difficult, since any significant test points to multiple potential culprits. Second, such composite null hypotheses can have lower power than univariate tests when only a small proportion of the phenotypes are associated with the tested genetic variant. This is a simple problem of dilution; a small number of true associations mixed with a large number of null phenotypes will reduce power. For these and other reasons the standard univariate test is often preferred to large multivariate analyses, although multivariate analyses are now considered when the number of phenotypes collected is small^9,13^.

## Have your cake and eat it

The objective of this work is to develop a method that keeps the resolution of univariate analysis when testing for association between an outcome *Y* and candidate predictor *X*, but takes advantage of other available covariates ***C*** = (*C*_1_, *C*_2_,… *C_m_*) to increase power. A first step toward this aim is to consider the inclusion of covariates correlated with the outcome in a standard regression framework. This may increase the signal-to-noise ratio between the outcome and the candidate predictor when testing: *Y* = *X* + ***C***. The selection of which covariates Q are relevant to a specific association test is usually based on causal assumptions^14–17^. Putting aside the estimation of indirect and direct effects^18^ of *X* on *Y*, epidemiologists and statisticians recommend the inclusion of two types of covariates: those that are potential causal factors of the outcome and independent of *X*, and those that may confound the association signal between *X* and *Y*, i.e. variables such as principal component (PC) of covariates that capture undesired structure in the data that can lead to false associations^19^. All other variables that vary with the outcome because of shared risk factors are usually ignored. However, those variables carry potentially interesting information about the outcome, and more precisely about the risk factors of the outcome. Because of their shared dependencies they can be used as proxies for risk factors of the outcome. As such, they can be incorporated in ***C*** to improve the detection of associations between *X* and *Y*. However, as we discuss further, when these variables depend on the predictor *X*, using them as covariates can lead to both false positive and false negative results depending on the underlying causal structure of the data.

The presence of interdependent explanatory variables, also known as multicollinearity^20^, can induce bias in the estimation of the predictor’s effect on the outcome. We recently discussed this issue in the context of genome-wide association studies that adjusted for heritable covariates^21–23^. To illustrate this *collider bias*, take first the simple case of two independent covariates *U*_1_ and *U*_2_ that are true risk factors of *Y*. When testing for association between *X* and *Y*, adjusting for *U*_1_ and *U*_2_ can increase power, because the residual variance of *Y* after the adjustment is smaller while the effect of *X* is unchanged, i.e. in **Figure 1a** the ratio of the outcome variance explained by *X* over the residual variance is larger after removing the effect of and *U*_2_. However, in practice, true risk factors of the outcome are rarely known. Consider instead the more realistic scenario where and *U*_2_ are unknown but a covariate *C*, which also depends on those risk factors, has been measured. Because of their shared etiology, *Y* and *C* display positive correlation, and when *X* is not associated with *C*, adjusting *Y* for *C* increases power to detect *(Y,X)* associations (**Fig. 1b**). Problems arise when *C* is associated with *X*. In this case adjusting *Y* for *C* biases the estimation of the effect of *X* on *Y*, decreasing power when the effect of *X* is concordant between *C* and *Y* (**Fig. 1c**), and inducing false signal when the effect is discordant (in opposite direction or when *X* is not associated with *Y*, **Fig. 1d**). These issues occur in **Figure 1c–d** because when *C* is included in the model, *U*_1_ and *U*_2_ become confounding variables between *X* and *Y* according to the causal graph.

**Figure 1.**
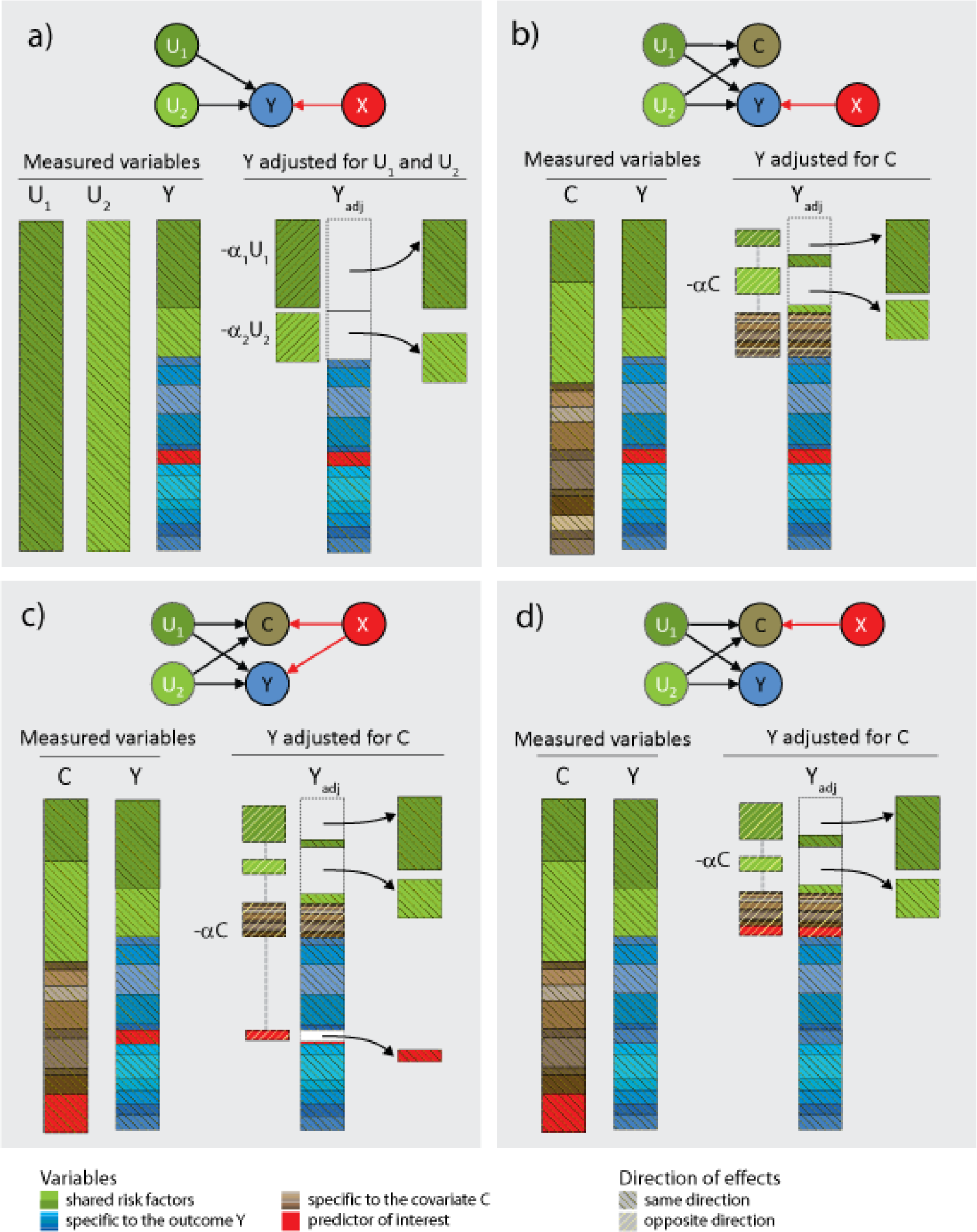
Variance components of adjusted variables. We illustrate the components of the variance of an outcome *Y* before and after adjusting for other variables. The predictor of interest, *X*, is displayed in red. In (a), the adjusting variables (*U*_1_ and *U*_2_) are true causal factors that have direct effects on *Y*, therefore adjusting *Y* for *U*_1_ and *U*_2_ reduce the variance of *Y*. In (b) the true factors are not measured but a variable *C* influenced by *U*_1_ and *U*_2_ is measured. Adjusting *Y* for *C* again reduces the residual variance of *Y*, but also introduces in the residual of *Y* a component of the variance specific to C. In (c) the covariate shares factors with *Y*, as in the previous scenario, but is also influenced by *X*. When the effect of *X* on *C* is concordant with the effect of *X* on *Y* (e.g. positive correlation between *C* and *Y*, and effect of *X* on *Y* and *C* in the same direction) this can induce loss in power, as the adjustment for *C* decreases the contribution of *X* to the residual of *Y*. In (d) *Y* is not associated with the predictor and adjusting for *C* can induce false association signal by introducing some effect of *X* in the residual of *Y*.

The same principles apply for any number of variables correlated with the outcome provided the sample size is large enough such that the effect of all covariates can be estimated in a multiple regression^2,4^. When none of the covariates depend on the predictor (**Fig. 1a–b**), their inclusion in a regression can reduce the variance of the outcome without confounding, leading to increased statistical power while maintaining the correct null distribution. This gain in power can be easily translated in terms of sample size increase. The noncentrality parameter (*ncp*) of the standard univariate test between *X* and *Y* equals 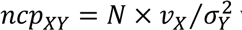 where *N*, *v*_x_ and 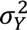 are the sample size, the variance of the outcome explained by the predictor, and the total variance of the outcome respectively. When reducing 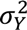 by a factor γ through covariate adjustment, and assuming the effect of *X* on *Y* is small, *ncp_XY_* can be approximated by 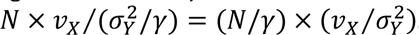. For example, when the covariates explain 30% of the variance of *Y*, the power of the adjusted test is equivalent to analyzing approximately a 1.4 fold larger sample size (as compared to the unadjusted test). When covariates explain 80% of the phenotypic variance-as discuss further, a realistic proportion in some genetic datasets-the power gain is equivalent to a 5 fold increases in sample size (**Fig. 2a**).

**Figure 2.**
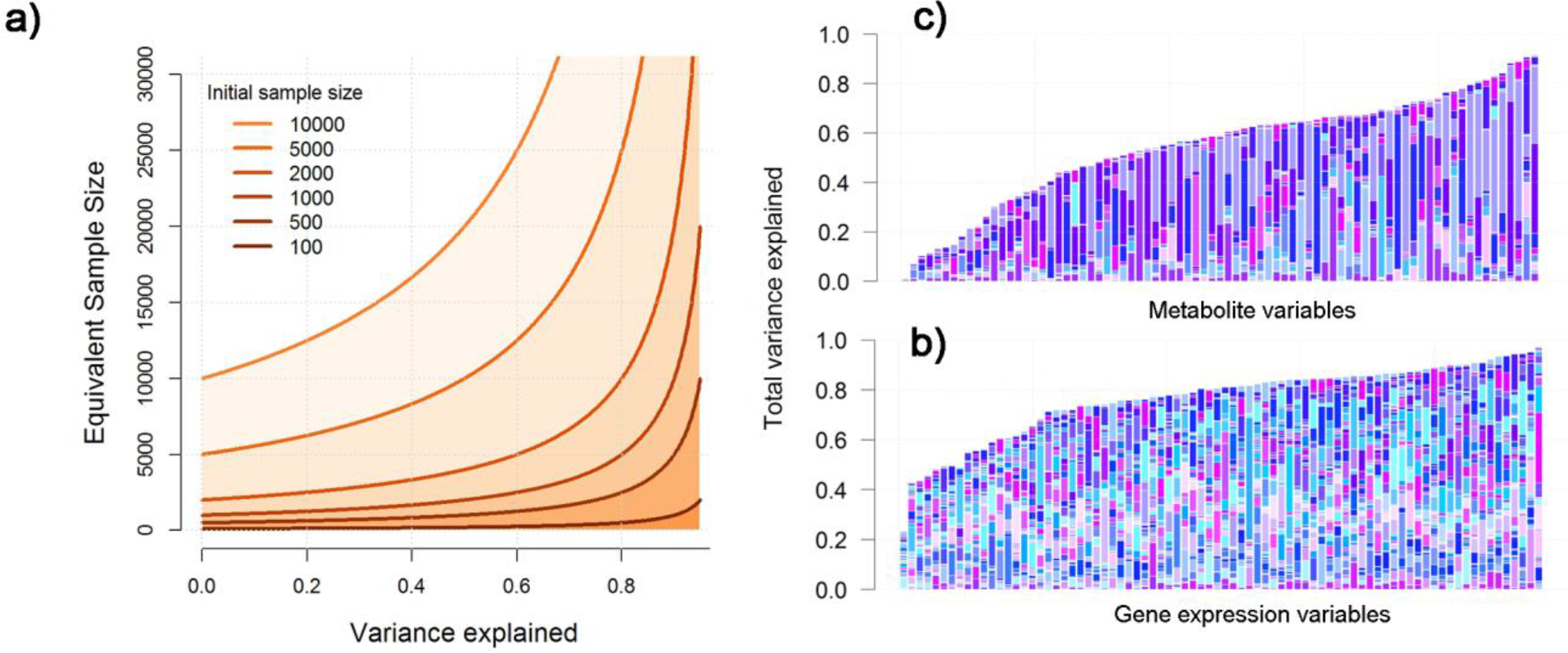
Variance components of adjusted variables. Equivalent increase in sample size as a function of the variance of the outcome explained by covariates assuming initial sample sizes ranging from 100 to 10,000 (a). Distribution of variance explained by other variables for the 79 metabolites from the PANSCAN study (b), and a random sub-sample of expression abundance estimates from 79 genes in the gEUVADIS study (c). The relative contribution of covariates to the total variance explained is illustrated with different sets of colors for each bar.

## Separating the wheat from the chaff

The central problem that must be solved is how to intelligently select a subset of the available covariates to optimize power while preventing induction of false positive or false negative associations. To do this, all covariates associated with the outcome should be included except those also associated with the predictor. A naïve solution would consist in filtering out covariates based on a *p*-value threshold from the association test between those covariates and the predictor considered. However, unless the sample size is infinitely large, some associations will be missed and unwanted covariates will be included. Furthermore, because a number of the covariates will be associated with the predictor by chance, the overall distribution of *p*-values from the covariate-adjusted test can be inflated, again potentially inducing false association signals (**Supplementary Fig. 1**). The underlying problem with *p*-value based filtering is that *p*-values are used to reject the null hypothesis in favor of the alternate. In our case the objective is to reject those covariates under the alternative hypothesis. Therefore, instead of using *p*-values to filter covariates, we developed a computationally efficient heuristic based on equivalence testing to improve the filtering of covariates while controlling the type I and type II error rate. Because the selected covariates will change for each predictor/outcome pair, we named our approach the *Musical Chair (MC)* algorithm (**Supplementary Fig. 2**).

Consider 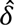, the estimated regression coefficient between *X* and *C*. The *p*-value based filtering can be transposed into an unconditional filtering on 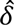. Under the null (δ = 0), 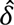 is normally distributed with mean 0 and variance 1/*N*, where *N* is the sample size. **Figure 3a** shows 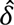 inclusion area for a p-value threshold of 5%-i.e. if 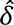 is outside the inclusion area, the covariate *C* is filtered out. Now consider 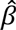, the estimated marginal effect of the predictor *X* on the outcome *Y* (not adjusted for *C*). Using 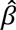 along with 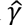 the estimated effect between *C* and *Y*, we can derive the conditional distribution of 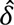, under a complete null model (β = 0 and γ = 0) (**Fig. 3b–c**, and **Supplementary Note**). The MC approach uses a joint inclusion area that combines the conditional and unconditional of distribution of 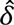. To prevent bias while maintaining power, the size of the inclusion region is determined by the strength of the collinearity between *X*, *Y*, and *C*.

**Figure 3.**
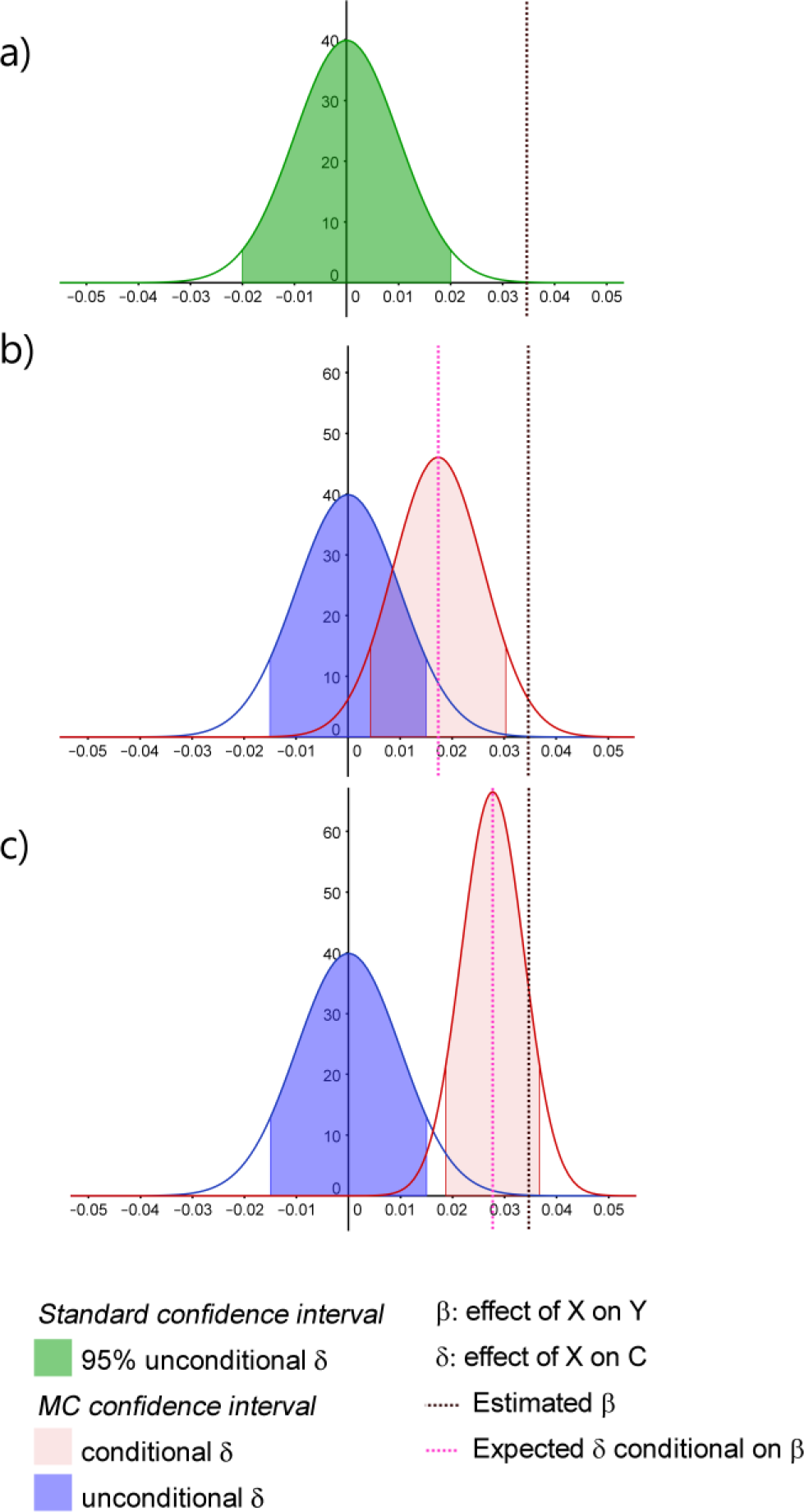
Conditional and unconditional distribution. Example of selection interval based on the distribution of 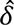, the estimated effect between the predictor *X* and the covariate *C* under the null hypothesis of no association between *X* and *C* (γ = 0) and no association between *X* and the outcome *Y* (β = 0). (a) presents the standard 95% confidence interval (green area) corresponding to p-value <0.05 unconditional on 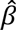. (b) and (c) show the composite interval of derived by the MC approach that merges and weights the expected unconditional (blue area) and conditional (pink area) distribution of 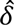 while considering a correlation between *Y* and *C* of 0.5 (b) and 0.8 (c). Plots were drawn assuming all variables are standardized, using a sample size of 10,000, an overall variance of *Y* explained of 0.7, 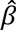 = 0.035 and a multivariate test of association between all covariates and *Y* with a *p*-value (*p*_Mul_) of 0.3.

More formally, for an outcome *Y*, a predictor *X*, and *m* candidate covariates *C* =(*C*_1_, *C*_2_,… *C*_m_), the MC algorithm uses four features to select covariates to be included in the model and perform a statistical test: i) *p*_MUL_, the *p*-value for the overall association between all *C*_l=1…*m*_, and *X*; ii) 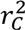 the amount of total outcome variance explained by the candidate covariates; iii) 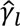 the estimated effect of each *C*_*l*∈1…*m*_ on *Y*; and iv)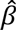 the marginal unadjusted estimated effect of *X* on *Y*. The first three features are used to define the stringency of the filtering (i.e. the size of the inclusion region). When *p*_MUL_ is very significant, the inclusion region is smaller reflecting the likelihood of the presence of undesired covariates. Similarly, when or 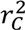 or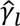 are large, the inclusion region is smaller because of potential bias^21^ (see e.g. **Fig. 3b**–**c**). The fourth feature,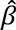, is used to make inference on the expected null distribution of 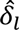, the regression coefficient between *X* and the covariate *C*_1_. It leverages the correlation between 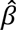 and 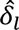 under a complete null model (β = 0 and δ = 0). These features are combined to derive a confidence interval Δ_l_ for each 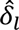, which determines whether a covariate can be safely included in the model.

Finally, when the dataset analyzed includes many covariates with substantial pairwise correlation (e.g. >0.2), the estimated effects of each covariate on *Y* obtained from a multivariate linear model,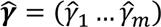, can vary substantially depending on which covariate is included in the model because of collinearity. This can be an issue because potential bias depends directly on 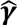. To address this point we implemented the selection of covariates described above into an iterative backward elimination where 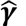 terms are re-estimated each time a candidate covariate is excluded. The complete details of the algorithm are presented in the ***online Methods*** section below.

## Simulated data analysis

We first assessed the performances of the proposed method through a simulation study in which we generated series of multi-phenotype datasets over an extensive range of parameter settings (see ***online Methods*** and **Supplementary Note**). Each dataset included *N* individuals genotyped at a single nucleotide polymorphism (SNP) with minor allele frequency (MAF) drawn uniformly in [0.1, 0.5], a phenotype *Y*, and *m* =[10,40,80] correlated covariates ***C*** = (*C*_1_, *C*_2_,… *C_m_*). Under the null, the SNP did not contribute to the phenotype and under the alternate the SNP contributed to the phenotype under an additive model. In some datasets, the SNP also contributed to a fraction *π* =[0%, 15%, 35%] of the covariates. These are the covariates, which we wish to identify and filter out of the regression. We considered sample sizes *N* of 300, 2,000 and 6,000, we varied 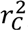, the variance of *Y* explained by ***C***, from 25% to 75%, and we increased the effect of the predictor on *Y* and ***C***, when relevant, so that it would corresponds to almost undetectable effects (i.e. median *χ*^2^ = 3) to relatively large effects (i.e. median *χ*^2^ = 20). For each choice of parameters we generated 10,000 replicates and performed four association tests: (unadjusted) linear regression (LR), linear regression with covariates included based on *p*-value filtering at an α threshold of 0.1 (FT), the MC algorithm (MC), and an oracle method that includes only the covariates not associated with the SNP (OPT), thus being the optimal test in regards of our goal. Crossing the different parameters, we considered a total of 351 scenarios which detailed results are presented in **Supplementary Figures 11-37**.

To comprehensively summarize the performances of the different tests across these many scenarios, we randomly sampled subsets of the simulations to mimic real datasets while focusing on a sample size of 2,000 individuals and a total of 100,000 SNPs tested. For null models we assumed that 70% of the genotypes would be under the complete null (not associated with any covariate, π = 0), while 20% would be associated with a small proportion of the covariates (π = 0.15) and the remaining 5% would be highly pleiotropic (π = 0.35). The results from this analysis are presented in **Figure 4**. Overall the MC approach outperforms the other methods (except OPT), being more powerful than LR with an average of two-fold increase in detection rate, and with dramatically lower false positives than FT (FT showed a genomic inflation factor^25^, λ_GC_ of 1.18, 1.17 and 1.19 when simulating 10, 40 and 80 covariates, respectively). However, in the extreme case when the number of candidate covariates is small (e.g. ≤10), they are highly correlated to the outcome (e.g. 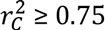), and the SNP is highly pleiotropic but has small effects, we observed a few outliers in the *p*-value distribution (e.g. **Supplementary Figs. 13,16,19**). Also, when the sample size was low compared to the number of covariate (e.g. *N* = 300, and *m*=10), we observed small deflation of the *p*-values under the null (**e.g. Supplementary Figs. 29-31**). We did not considered the strategies which consists in including all *C*_*l*=… *m*_ variables as covariates without any filtering on predictor-covariate association or the so-called reverse regression (which consists in using the predictor as the outcome^26^), as both approaches lead to substantial type I error rate (see **Supplementary Fig. 3**).

**Figure 4.**
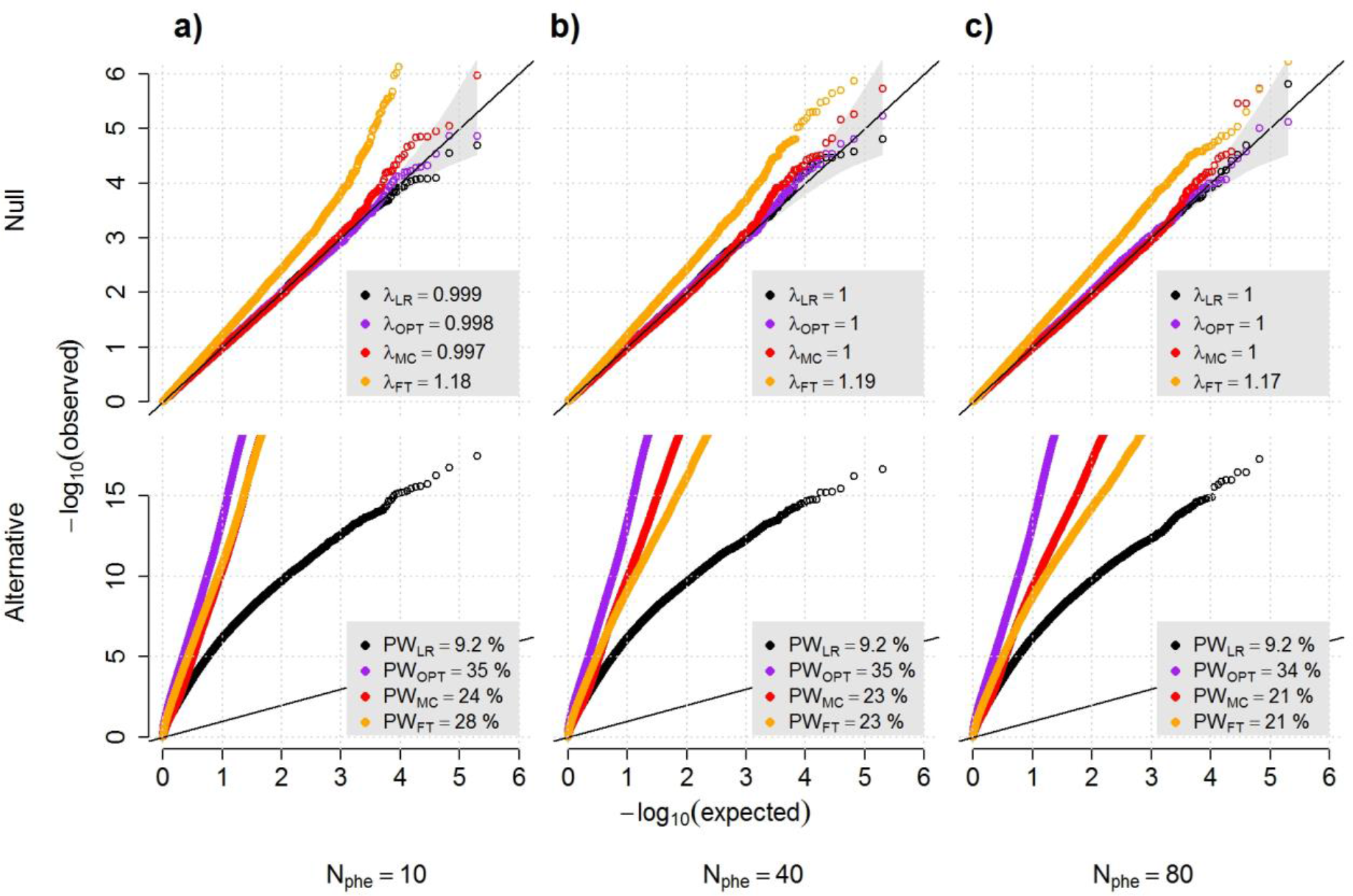
Power and robustness. We simulated series of 100,000 datasets including 10 (a), 40 (b) and 80 (c) outcomes under a null model (upper panels), where a predictor of interest is not associated with a primary outcome but is associated with either 0%, 15% or 35% of the other outcomes with probability 0.75, 0.2 and 0.05 respectively, and under the alternative (lower panels), where the predictor is associated with the primary outcome only. The variance of the primary outcome that can be explained by the other outcomes was randomly chosen in [25%, 50%, 75%] with equal probability. In each replicate we applied four tests of association between the primary outcome and the predictor: a standard marginal univariate test (LR); the optimally adjusted test (OPT) that includes as covariates only the outcomes not associated with the predictor; the MC test (MC); and a univariate test that include as covariate all outcomes with a *p*-value for association with the predictor above 0.1 (FT). For the null models we derived the genomic inflation factor *λ_GC_*, while for the alternative model we estimated power at an α threshold of 5x10^−7^, to correct for 100,000 tests.

## Real data analysis

We first analyzed a set of 79 metabolites measured in 1192 individuals genotyped at 668 candidate single nucleotide polymorphisms (SNPs). We derived the correlation structure between these metabolites (**Fig. 2b** and **Supplementary Fig. 4**)^5^ and estimated the maximum gain in power that can be achieved by our approach in these data. The proportion of variance of each metabolite explained by the other metabolites varied between 1% and 91% (**Fig. 2b**). This proportion is higher than 50% for two thirds of the metabolites, meaning that for all those variables, one can potentially achieve a gain in power equivalent to a two-fold increase in sample size. More interestingly, for 10% of the metabolites, other variables explain over 80% of the variance, corresponding to a maximum five-fold increase in sample size. In such cases, predictors explaining a very small amount of a metabolite’s variation (e.g. <1%) can moved from undetectable (power<1%) to fully detectable (power>80%). We performed a systematic screening for association between each SNP and each metabolite, using both a standard univariate linear regression adjusting for potential confounding factors and using the MC approach to identify additional covariates. Overall, both tests showed correct *λ_GC_* (**Supplementary Fig. 5**). We focused on associations significant after Bonferroni correction (*P* < 9.5×10^−7^ corresponding to a correction for the 52,772 tests performed). The standard unadjusted approach (LR) detected 5 significant associations. In comparison, the MC approach identifies 10 associated SNPs (**Table 1**), including four of the five associations identified by LR. In most cases the *p*-value of our approach was dramatically lower (e.g. 1000 fold smaller for the rs780094-alanine association). Comparing these results to four previous independent GWAS metabolite scans of larger sample size (*N* equal 8,330, 7,824, 2,820, and 2,076 for Finnish^27^, KORA+TwinsUK^6,28^, and FHS^29^, respectively), we found that all metabolite/gene associations only identified by the MC approach have been previously identified (**Supplementary Table 1**). These positive controls confirmed the power of the proposed MC approach, highlighting its ability to identify variants with much smaller sample size. Interestingly, the only association identified by the unadjusted analysis (lactose and *GC*, *P*=6.1×10^−7^) and not confirmed by the MC approach (*P*=6.3×10^−6^) was also the only one not previously reported in previous larger studies, highlighting the ability of the proposed MC approach to improve not only power but also type II error rate.

**Table 1.**
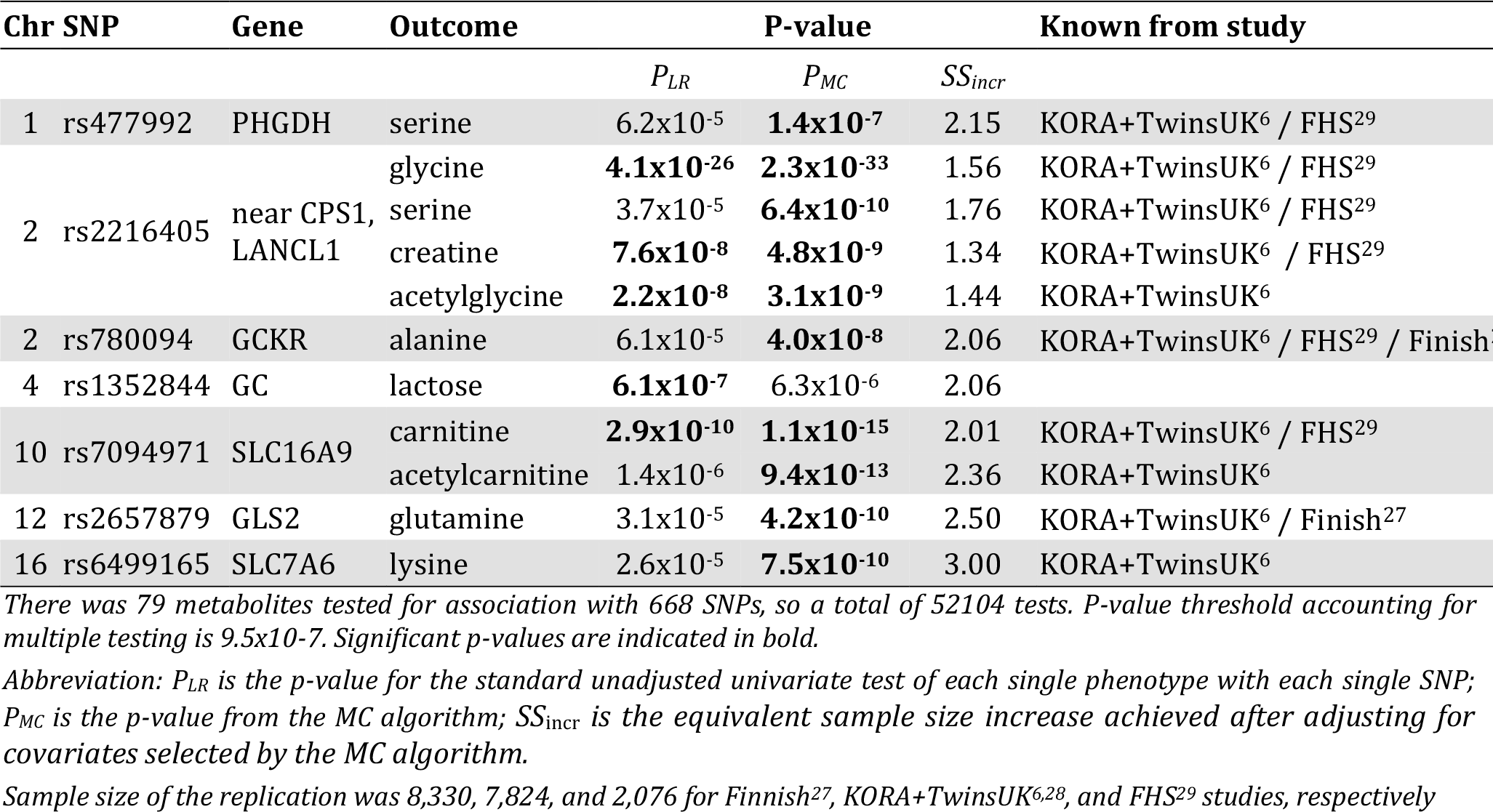
Identified signals from the association test between 79 metabolites and 668 candidate SNPs.

We then considered genome-wide *cis*-eQTL mapping in RNA-seq data from the gEUVADIS study. Gene expression is a particularly compelling benchmark, as the current standard analyses already use an adjustment strategy to account for hidden factors in *trans* and *cis* eQTL GWAS^30^–^33^. Here we used the PEER approach^30^ to derive those hidden factors, as the method has been applied in one of the major recent cis-eQTL screenings in the gEUVADIS data^34^. After stringent quality control the data included 375 individuals of European ancestry with expression estimated on 13,484 genes, of which 11,694 had at least one SNP with a MAF ≥ 5%. We observed that expressions levels between genes were highly correlated (**Fig. 2c**), an ideal scenario for MC. We first performed a standard cis-eQTL screening using linear regression (LR), testing each SNP within 100kb of each available gene for association with overall RNA level while adjusting for 10 PEER cofactors, for a total of ~1,3 million tests. Then, we applied MC to identify for each test, which other gene’s RNA levels could be used as covariates on top of the PEER factors. As shown in **Supplementary Figure 6**, both LR and MC showed large number of highly significant association. For comparison purposes we plotted in **Figure 5** the most significant SNP per gene obtained with the standard approach against those obtained with MC. As shown in this figure, 2,725 genes had a least one SNP significant with both methods, and 56 genes were identified by the standard approach only. Conversely 657 genes were found only with the MC approach, corresponding to a 24% increase in detection of *cis*-eQTLs. This indicates that by being gene/SNP specific, the MC approach is able to recover substantial additional variance, allowing for increased power. We also performed quasi-null experiment where we tested for cis-effect using random SNPs from the genome. We observed a small inflation (λ_LR_=1.01, and λ_MC_=1.05, **Supplementary Figs. 7-8**). However even after correcting *p*-value of the former analysis for this potential bias the improvement in detection for MC remained above 22%.

**Figure 5.**
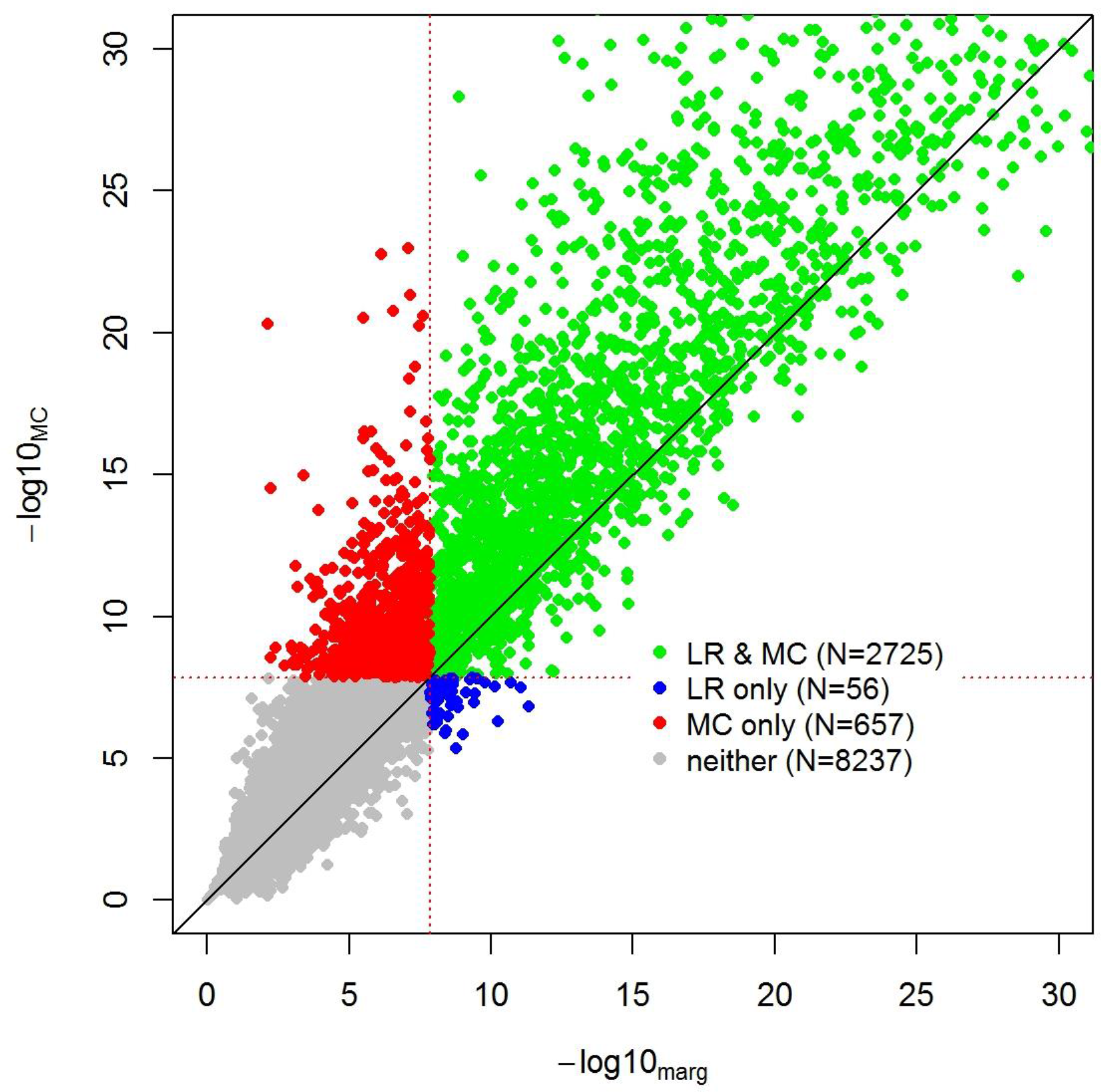
Analysis of the gEUVADIS data. We performed a genome-wide *cis*-eQTL mapping of 11,694 genes in 375 individuals from the gEUVADIS study. Analysis was performed using standard linear regression (LR) and the *Musical Chair* (MC) approach. Both consisted in running a linear regression adjusted for 10 PEER factors, while the MC analysis also included 0 to 50 additional covariates per SNP/gene pair tested. We compared the-log_10_(*p*-value) of the most significant SNP per gene obtained by each approach. For illustration purposes we shrunk the plots at-log_10_(*p*-value)=30. We considered a stringent significance threshold of 1.4×10^−8^ to account for the approximately 3.5millions test and derived the number of gene showing at least one cis-eQTL with LR only (blue), MC only (red), both approaches (green) or neither (grey).

## Discussion

Growing collections of high-dimensional data across myriad fields, driven in part by the “big data revolution” and the *Precision Medicine Initiative*, offer the potential to gain new insights and solve open problems. However, when mining for associations between collected variables, identifying signals within the noise remains challenging. While univariate analysis offers precision, it fails to leverage the correlation structure between variables. Conversely multivariate methods have increased power at the cost of decreased precision. We demonstrated in both simulated and real data that the proposed method, *Musical Chairs*, maintains the precision of univariate analysis, but can still exploit global data structures to increase power. Indeed, in the data sets examined in this study we observed up to a 3-fold increase in effective sample size in both the gene expression and metabolites data (**Supplementary Figure 9**) thanks to the inclusion of relevant covariates. Moreover, results from other ongoing applications of our approach to other real datasets show promise. In particular, we recently used MC to screen for association between gut microbiome and genetic variants in individuals with inflammatory bowel disease. The MC approach allowed for the identification of an association between a risk score for *NOD2* and *F,prausnitzii* which was missed by the standard approach.^35^ This result, in agreement with recent functional studies^36,37^was further confirmed in a replication dataset using the standard (unadjusted) approach.

*Musical Chairs* can be potentially applied to any type of data, however it is particularly well suited to the analysis of human genomic data for several reasons. First, the genetic architecture of human phenotypes likely follows a polygenic model with many genetic variants of small effect size that are difficult to detect using standard approaches^38^. Second, many correlated phenotypes share genetic and environmental variance without complete genetic overlap^39^. Each single phenotype from a multi-phenotype dataset depends on a mixture of shared and phenotype-specific risk factors, and the aforementioned principle can be applied. Third, the underlying structure of the genomic data is relatively well understood with an extensive literature on the causal pathway from genotypes to phenotypes through direct and indirect effects on RNA, protein and metabolites^40^ (**Supplementary Fig. 10** and **Supplementary Note**). Finally, when the predictors of interests are genetic variants, e.g. single nucleotide polymorphisms (SNPs), there is little concern regarding potential confounding factors. The only well-established confounder of genetic data is population structure and this can be easily addressed using standard approaches^19^. For other types of data, application should be considered on a case-by-case basis. In particular, when the underlying structure of the data is unknown the risk for introducing bias is higher, especially when the many variables have causal relationships. Second, confounding factors will in general match covariate’s criteria for exclusion as they are, by definition, correlated with both the outcome and the predictor. These covariates should indeed remain in the model and our approach allows for their inclusion. However, as for any large scale screening using standard approach, manually defining confounding factors for each predictor/outcome pair can be a daunting task. Moreover, confounding factors might not always be well known.

Several other groups have considered the problem of association testing in highdimensional data. In genetics, multivariate linear mixed models (mvLMMs) have demonstrated both precision and increases in power when correlated phenotypes are tested jointly^9^. However, mvLMMs are only exploiting the genetic similarity of phenotypes and are not computationally efficient enough to handle dozens of phenotypes jointly (e.g. would be limited to the analysis of 2 to 10 phenotypes^10^) let alone hundreds. MC leverages both genetics and environmental correlations and can be easily adapted to hundreds or thousands of phenotypes as we demonstrated here. It is also worth noting that substantial work has been published on how to account for hidden technical artifact in *trans* and *cis* eQTL GWAS^30–33^. While adjusting for principal components of expression has been the most common approach^33^ and is still commonly used, other, more complex methods have been proposed, including SVA^31^ and PEER^30^ (which we used in parallel with MC in the gEUVADIS analysis). Though presented from a different perspective, these methods aim at recovering what would be *U*_1_ and *U*_2_ in **Figure 1**. One advantage of these methods is that they reconstruct hidden factors only once for all of the outcomes data, thus being more computationally efficient. However, by not being specific they can (i) induce false signal if genetic effects happen to be captured by these factors, and (ii) be suboptimal, as they assume a limited number of shared risk factors while our approach does not make such assumption and optimizes the test for each predictor-outcome pair. Indeed the gEUVADIS analysis showed a 24% increase in the detection of eQTL when applied on top of PEER.

There are several caveats to our approach. First, the proposed heuristic is conservative by design in order to avoid false association signal and so all the available power gain is not achieved. Second, while all simulations we performed show strong robustness of our approach, it remains a heuristic, and the validity of the proposed approach cannot be guaranteed. We are currently examining alternatives for excluding covariates, such as structural equation modelling, which more directly assess causal relationships at the expense of computational efficiency^41^. Ultimately we recommend external replication to validate results as is standard in genetic studies. Third, MC is more computationally intensive than methods such as PCA or PEER which derived hidden factors for all data at once. However, as we demonstrated here, this is the cost for improved statistical power. Still, we are actively working on updates of the algorithm to improve computational efficiency. Fourth, the method assumes that the variables are measured and available on all samples and we intend to explore the handling and imputation of missing phenotypes in future work. Fifth, while principles we leveraged are likely applicable to categorical and binary outcome (see e.g. ^42^ for logistic regression), as of now, our algorithm is only applicable to continuous outcomes. There are also other additional improvements not specific to MC that might be worth exploring in future works. In particular, when multiple phenotypes are considered as outcomes then a multiple test correction penalty must be selected to account for all tests across all phenotypes. In this work we applied a Bonferroni correction, not accounting for the correlation between outcomes. This is a very conservative correction and more powerful approaches are possible.

Big genomic data have the potential to answer important biological questions and improve public health. However those data come along with great methodological challenges. Many questions, such as improving risk prediction or inferring causal relationship, rely in particular on our ability to identify association between variables. In this study we provide a comprehensive overview of how leveraging shared variance between variables can be used to fulfill this goal. Building on this principle we developed the *Musical Chair* algorithm, an innovative approach which can dramatically increase statistical power to detect weak association.

## Online Methods

### Principle

Consider two variables *Y* and *C* that are collected on the same set of individuals, which we would like to examine in an association study. If *Y* and *C* are correlated then they either influence each other or they share sets of risk factors. In the latter case, it means that the variations of the shared risk factors are captured by both variables. Therefore, *C* can be a proxy for a risk factor or a (eventually, a complex) combination of risk factors of *Y* and conversely, *Y* can be a proxy for risk factors of *C*. When a predictor *X* is not part of these shared risk factors and is for example associated with *Y* only, it implies that including *C* as a covariate in the association test between *X* and *Y* can increase power since part of the variance of *Y* not explained by *X* will be removed. Consider the simple additive linear model in which *Y* is generated according to:

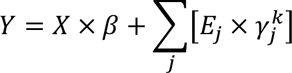

where *X* is a measured risk factor of *Y* and *E*_*J*=1…*k*_ are unmeasured risk factors of *Y*. The expected test statistic when testing the association between the normalized predictor *X* and *Y* is *cor*(*X, Y*)^2^ × *N*, where *N* is the number of individuals in the study. This correlation is a function of β and the variances of *X* and *Y*. If it were possible to create a new adjusted outcome:

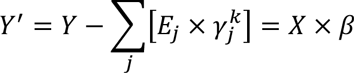

Then the correlation *cor*(*X*, *Y*’)^2^ × *N* = *N*, and this would be optimally powered. If another variable *C* collected in the study is correlated with *Y*, then it might share causal risk factor *X* and/or some of the *E*_j =1…k_. We can include this variable as a covariate in the regression when testing for association between *X* and *Y*. If *X* is not associated with *C*, then this is effectively removing elements of *E* that influence *Y* and thereby increasing the power of the association test.

The issue with the application of this principle is that if *X* is associated with *C*, then including it as a covariate in the regression will potentially decrease the test statistic since elements of *X* will be removed from *Y*. Even worse, if *X* is associated with *C* only, then including *C* as a covariate can induce a false association signal. The objective of our approach is to remove from *Y* variance explained by factors not correlated with *X* in order improve the study’s power.

### The heuristic

We develop a heuristic to select relevant covariates when testing for association between a predictor *X* and an outcome *Y*. For a set of candidate covariates ***C*** = (*C*_1_, *C*_2_,  *C*_m_), the filtering is applied on 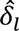 and *p_l_*, the estimated marginal effect of the predictor *X* on Q and its associated *p*-value, respectively. It uses four major features: i)*p*_MU_, the *p*-value for the multivariate test of all *C*_l=…m_ and *X*, which is estimated using a standard multivariate approach (a MANOVA in the present application); ii)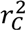 the total amount of variance of *Y* explained by the ***C***; iii)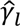 the estimated effect of each *C*_*l*∈1…m_ on *Y*; and iv)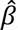, the estimated effect of *X* on *Y* the marginal model *Y*~α + β*X*.

Filtering is applied in two steps using the aforementioned features and additional parameters describe thereafter. Step 1 is an iterative procedure focusing on *p*_MUL_. It consists in removing potential covariates until *p*_MUL.s_ reaches *t*_MUL_, a *p*-value threshold. This step is effective at removing combination of covariates with strong to moderate effects, but will potentially leave weakly associated covariates. Step 2 is also iterative and uses covariates pre-selected at step 1. It consists in deriving two confidence intervals Δ_I.cond_ and Δ_i.un_, for the expected distribution of 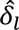 conditional on 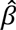 under a complete null model (δ_*l*_ = 0 and α = 0), and the unconditional distribution of 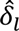, respectively. The unconditional distribution of 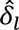 can be approximated as 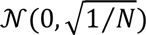, while the conditional distribution equals 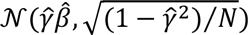, where 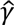 is the estimated correlation between *Y* and *C* (see **Supplementary Notes**). The final inclusion area for each 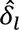 is then defined as the union of Δ_*l.cond*_ and Δ_*l.un*_ after applying *ad hoc* weighting functions. This includes first a stringency weight w_ST_ that combines the aforementioned indicators of potential bias. The second component consists in two semi-linear threshold functions *f*_c_ and *f*_u_ that balance the importance of the two inclusion areas of each *c*_*l*_ (i.e. Δ_*l.cond*_ and Δ_*l.un*_, respectively) in order to reflect the probability of being under a complete null model, and to limit them to be no larger than the 95% confidence interval (CI) of 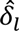. More specifically, when 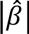 is small, the two intervals (Δ._*l.cond*_ and Δ_*l.un*_) are giving the same weights, however as 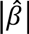 increases, the likelihood of the true β being null decreases and the conditional interval, Δ_*l.cond*_ is shrunk to zero. In practice we used simple linear functions with a tipping point that corresponds to a situation where the 95% *CI* of the observed 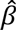 and 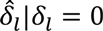 stop overlapping. The former *CI* approximately equals 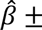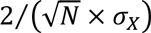 where σ_x_ is the standard deviation of *X*, while the later equals 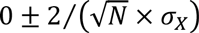. Expressed as chi-squared this tipping point corresponds to 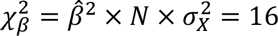.

The proposed multi-step algorithm is defined as follows:

For each predictor *X* and *Y*

1. Univariate association

1. 1.1. Standardized all variables (*Y, X, **C***) to have mean 0 and variance 1
2. 1.2. Initialize *L* = 1…*m*, the list of selected covariates, with all available covariates
3. 1.3. Derive for each *l*∈ *L*, 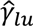 and 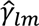 the marginal effect estimates from the univariate regression *Y*~*C*_*l*_=_*l*…*m*_, and multivariate model *Y*~***C***, respectively.
2. Filter 1: multivariate

1. 2.1. Perform a marginal association test between *X* and each *C_t_*=_*l*…*m*_ Derive all 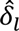 and *p*_*l*_ from *Y*~*δ*_0_ + δ_*l*_×*X*
2. 2.2. Set*p*_MUL_=1
3. 2.3. While *P_MUL_* < *t*_*MUL*_
  2.3.1. Derive *P*_MUL_ from *C_L_*~*X* using a multivariate test, where *C_L_* is the data matrix *C* including only *I* ∈ *L* covariates.
  2.3.2. Update *L* by removing the *C_l_* that match *p_*i* ∈ *L*_ = min(*p*_*l*∈*L*_) from the set of candidate covariates*

1. if L ≠ 0, filter2: univariate

1. 3.1. while L ≠ 0 and L_t+1_ ≠ L_t_
  1. 3.1.1. Update for each l∈ L 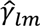 the effect estimates from the multivariate model *Y*~***C***_L_
  2. 3.1.2. Derive 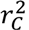 the variance of Y explained by ***C***_L_ from the model in 3.1.1
  3. 3.1.3. Derive for each *l* ∈ *L* the *ad hoc* stringency weight of the inclusion area 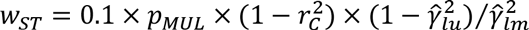
  4. 3.1.4. Derive the overall weights of the conditional and unconditional models using semi-linear threshold where 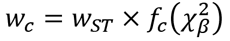 and 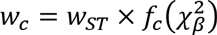 where 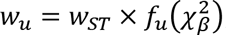

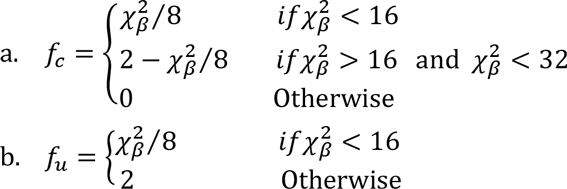
  5. 3.1.5. Derive the mean *μ_l.un_* = 0 and standard deviation 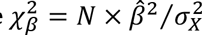 of the unconditional distribution of 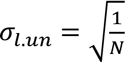 and the associated inclusion area:

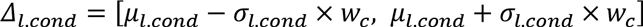
  6. 3.1.6. Derive the mean 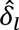 and standard deviation 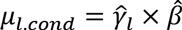 of the conditional null distribution of 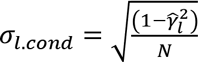, and the associated inclusion area:

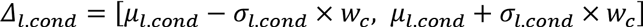
  7. 3.1.7. Update *L* by removing all *l* which 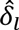 is not included in *Δ_l.cond_* ∪ *Δ_l.un_*
2. Perform the test of association between *X* and *Y*, while adjusting for the selected covariates

1. 4.1. Estimate 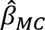 and derive the associated *p*-value from the multivariate model including all *l* ∈ *L* covariate from *Y*~ β_0_ + × X + ***β_L_*** × ***C_L_***

### Simulations

We simulated *Y* and *m* correlated phenotypes *Y*, ***C*** = (*C*_1_, *C*_2_,… *C_m_*) under a variety of genetic models to interrogate the properties of the proposed test. Genotypes *g* for each of *N* individuals were generated by summing two samples from a random binomial distribution with probability uniformly drawn in [0.1, 0.5] and then normalized to have mean 0 and variance 1. Under the alternate, the effect of the genotype on phenotype *Y* had effect size β, and effect size 0 under the null. In some simulation, the genotype was also associated to a fraction *π* of the *m* covariates with effect size drawn from [-*δ*, *δ*]. The remaining variance for each phenotype was drawn from a *m+1-dimensional* multivariate normal distribution, and represents the remaining genetic and environmental variance. The diagonal of the covariance matrix was specified as *1 minus* the effect of *g* (if relevant) such that the total variance of each phenotype had an expected value of
1. The off diagonal elements for each pair of phenotypes specifies the phenotypic covariance and was drawn from a normal distribution with mean 0 and variance *σ_c_*. In instances where this matrix was not positive definite we used the Higham algorithm^43^ to find the closest positive definite matrix. For each null model we derived the genomic inflation factor^25^ *λ_GC_*, while for the alternative model we estimated power at an *α* threshold of 5×10^−7^, to account for the 100,000 tests performed.

### The metabolite data

Circulating metabolites were profiled by liquid chromatography-tandem mass spectrometry (LC-MS) in prediagnostic plasma from 453 prospectively-identified pancreatic cancer cases and 898 controls. These subjects were drawn from four U.S. cohort studies: the Nurses Health Study (NHS), Health Professionals Follow-up Study (HPFS), Physicians Health Study (PHS) and Women’s Health Initiative (WHI). Two controls were matched to each case by year of birth, cohort, smoking status, fasting status at the time of blood collection, and month/year of blood collection. Metabolites were measured in the laboratory of Dr. Clary Clish at the Broad Institute using the methods described in Wang et al.^44^ and Townsend et al.^45^ A total of 133 known metabolites were measured; 50 were excluded from analysis because of poor reproducibility in samples with delayed processing (n=32), CV>25% (n=13), or undetectable levels for >10% subjects (n=5). The remaining 83 metabolites showed good reproducibility in technical replicates or after delayed processing.^45^ Among those, 79 had no missing data and were considered further for analysis. Additional details of these data have can be found here^46^. Genotypic data was also available for some of these participants. A subset of 645 individuals from NHS, HPFS and PHS had genome-wide genotypes data as part of PanScan study^47^. Among the remaining participants, 547 have been genotyped for 668 SNPs chosen to tag genes in the inflammation, vitamin D, and immune pathways. To maximize sample size we focused our analysis on these 668 SNPS which were therefore available in a total of 1,192 individuals. Insample minor allele frequency of these variants range from 1.1% to 50%. We first applied standard linear regression testing each SNP for association with each metabolite while adjusting for five potential confounding factors: pancreatic cancer case-control status, age at blood draw, fasting status, self-reported race, and gender. We then applied the MC approach while also including the five confounding factors as covariates.

### The gEUVADIS data

The gEUVADIS data^34^ consists of RNA-seq data for 464 lymphoblastoid cell line (LCL) samples from five populations in the 1000 genomes project. Of these, 375 are of European ancestry (CEU, FIN, GBR, TSI) and 89 are of African ancestry (YRI). In these analyses we considered only the European ancestry samples. Raw RNA-sequencing reads obtained from the European Nucleotide Archive were aligned to the transcriptome using UCSC annotations matching hg19 coordinates. RSEM (*RNA-Seq* by Expectation-Maximization)^48^ was used to estimate the abundances of each annotated isoform and total gene abundance is calculated as the sum of all isoform abundances normalized to one million total counts or transcripts per million (TPM). For each population, TPMs were log2 transform and median normalized to account for differences in sequencing depth in each sample. A total of 29,763 total genes were initially available. We removed those that appear to be duplicates or that had low expression value (defined as log2(TPM)<2 in all samples). After filtering, 13,484 genes remain. The genotype data was obtained from 1000 Genomes Project Phase 1 data set. We restricted the analysis to the SNPs with a MAF≥5% that were within ±50kB from the gene tested for cis-effect A total of 11,694 genes had at least one SNP that match these criteria.

When running the MC approach, we performed a pre-filtering of the candidate covariates. More specifically, for each gene analyzed-referred further as the *target* gene-we restrained the number of candidate covariates (i.e. gene other than the *target*) to be evaluated. First, we aimed at avoiding genes which expression is more likely to be associated with some of the SNPs tested because of a *cis*-effect, as such genes are more likely to induce false signal. Thus, all genes in close physically proximity with the target genes (≤1Mb) were excluded. Second, we aimed at reducing the number of candidate covariates (13,484 minus 1, *a priori*), as most of them are likely uninformative and also because our simulation showed that for small sample size, the MC approach would have reduced robustness if the number of candidate covariates is too large. To do so we performed an initial screening for association between the *target* and all others genes and used further the top 50 showing the strongest squared-correlation with the *target*.

Because of a dramatic number of true associations, the main cis-eQTL screening showed strong genomic inflation factors (Figure S6, *λ_LR_*=2.21, *λ_MC_*=2.30). Therefore, to assess the validity of the MC approach, we repeated the analyses above, but tested each gene’s expression with sets of SNPs chosen on a different chromosome, in order to preserve both the expression correlation and the SNPs correlation. This analysis almost corresponds to a null, although some *trans* effect might be captured in this experiment. The *λ_GC_* was slightly inflated (*X_LR_ =1.01* and *λ_MC_* =1.05, **Supplementary Fig. 7-8**) but did not display any strong outlier.

### Variance explained in multiple regressions

We plotted in Figure 2b–c the variance of a set of outcomes *Y=* (*Y_1_*,… *Y_K_*) that can be explained by covariates in the data-i.e. how much of the variance of *Y_i_* can be explained by *Y_j≠i_*. For illustration purposes we also approximate the individual contribution of each *Y_j≠i_* covariate. In brief, we standardized all variables and estimated 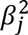, the proportion of variance of the outcome explained by each *Y_j≠i_* from the models *Y_i_*~***βY***_*j*#x2260;*i*_, and 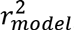, the total variance of *Y_i_* explained by all *Y_j≠i_* jointly, from the model *Y_i_*~*βY*_*j*#x2260;*i*_. Then, we derived *v_ij_* the relative contribution of each *Y_j≠i_* to the variance of *Y_i_* as follows:

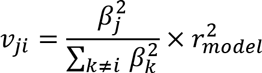

This is only an approximation of the real contribution of each variable, since the interdependence between the covariates implies instability of all estimates. Indeed adding or removing covariates often leads to changes of the *β_j_*.

## Author contributions

H.A. conceived the approach. N.Z. supervised the project. H.A., N.Z., B.V., C.P., D.S. and P.K. contributed substantially to improvement of the approach and the study design. J.Y. contributed to the quality control and analysis of the gEUVADIS data. B.W. collected the metabolites data and contributed to the quality control and analysis of the metabolites data. H.A. and N.Z. conceptualized and performed the simulation study. H.A. performed all real data analyses. H.A. and N.Z. wrote the manuscript.

